# The maternal genetic history of the Angolan Namib Desert: a key region for understanding the peopling of southern Africa

**DOI:** 10.1101/162230

**Authors:** Sandra Oliveira, Anne-Maria Fehn, Teresa Aço, Fernanda Lages, Magdalena Gayá-Vidal, Brigitte Pakendorf, Mark Stoneking, Jorge Rocha

**Affiliations:** CIBIO/InBIO: Research Centre in Biodiversity and Genetic Resources, University of Porto, Portugal; Departamento de Biologia, Faculdade de Ciências, Universidade do Porto, Portugal; Department of Linguistic and Cultural Evolution, MPI for the Science of Human History, Jena, Germany; Institute for African Studies, Goethe University, Frankfurt, Germany; Centro de Estudos do Deserto (CEDO), Namibe, Angola; ISCED/Huíla—Instituto Superior de Ciências da Educação, Lubango, Angola; Laboratoire Dynamique du Langage, UMR5596, CNRS & Univ Lyon, Lyon, France; Department of Evolutionary Genetics, MPIfor Evolutionary Anthropology, Leipzig, Germany

## Abstract

Southern Angola is a poorly studied region, inhabited by populations that have been associated with different migratory movements into southern Africa. Besides the long-standing presence of indigenous Kx’a-speaking foragers and the more recent arrival of Bantu-speaking pastoralists, ethnographic and linguistic studies have suggested that other pre-Bantu communities were also present in the Namib desert, including peripatetic groups like the Kwepe (formerly Kwadi speakers), Twa and Kwisi. Here we evaluate previous peopling hypotheses by analyzing the relationships between seven groups from the Namib desert (Kuvale, Himba, Tjimba, Kwisi, Twa, Kwepe) and Kunene Province (!Xun), based on newly collected linguistic data and 295 complete mtDNA genomes. We found that: i) all groups from the Namib desert have genealogically-consistent matriclanic systems that had a strong impact on their maternal genetic structure by enhancing genetic drift and population differentiation; ii) the dominant pastoral groups represented by the Kuvale and Himba were part of a Bantu proto-population that also included the ancestors of present-day Damara and Herero peoples from Namibia; iii) Tjimba are closely related to the Himba; iv) the Kwepe, Twa and Kwisi have a divergent Bantu-related mtDNA profile and probably stem from a single population that does not show clear signs of being a pre-Bantu indigenous group. Taken together, our results suggest that the maternal genetic structure of the different groups from the Namib desert is largely derived from endogamous Bantu peoples, and that their social stratification and different subsistence patterns are not indicative of remnant groups, but reflect Bantu-internal variation and ethnogenesis.

## INTRODUCTION

The high ethnic diversity of southwestern Angola, the importance of its pastoral culture, and the likely confluence of different migratory waves in its peopling provide a unique opportunity to explore the significance of different hypotheses about the population history of southern Africa. At present, it is generally accepted that the oldest population stratum in this vast region is represented by groups speaking languages that make extensive use of click consonants, which were previously lumped into a hypothetical “Khoisan” phylum (Greenberg 1963), but are now divided into three families: Kx’a, Tuu and Khoe-Kwadi (Güldemann and Fehn 2014). While Tuu and Kx’a-speaking peoples were historically hunter-gatherers, Khoe-Kwadi languages are spoken by both foraging and food-producing groups, with the Khoekhoe-speaking Nama representing one of the major pastoralist populations of southern Africa. Based on typological observations, it has been speculated that the Khoe-Kwadi languages might constitute a later arrival in the area, possibly linked to a migration of Later Stone Age pastoralists from East Africa, who moved into regions previously inhabited by Kx’a and Tuu speaking hunter-gatherers (Westphal 1963; Barnard 1992; Güldemann 2008). Although the presence in southern Africa of lactase persistence and Y-chromosome haplotypes that originated in eastern African pastoralists seems to support this hypothesis (Henn et al. 2008; Coelho et al. 2009; Breton et al. 2014; Macholdt et al. 2014), it is unclear whether these traits were dispersed by a massive immigration of Khoe-Kwadi speakers or were introduced through small scale movements leading to the diffusion of livestock and genetic variants across neighboring resident populations (Sadr 2015).

More recently, about 1,500 years ago, the human population landscape of southern Africa was further modified by the arrival of Bantu-speaking groups with subsistence economies that presently range from almost exclusive pastoralism to mixed farming systems (Russell et al. 2014). While the emergence of new combinations of genes, languages and modes of subsistence is an expected outcome of the confluence of different population strata, the prevailing views about the peopling of southern Africa favor the idea that the technological advantages and social dominance of the Bantu considerably restricted the direction and range of genetic and cultural exchange (Cashdan 1986). Consequently, a strong connection between foraging, low social status, the “Khoisan”languages and phenotypes including small stature and light skin was established, leaving anthropologists puzzled with foraging peoples physically more similar to other non-“Khoisan” African populations (Cashdan 1986; Barnard 1992). In this context, the origin of the dark-skinned foragers speaking Khoe-Kwadi languages, such as the Khwe from the Okavango region, or the Damara from Namibia, is often considered enigmatic and has been linked to a hypothetical stratum of pre-Bantu non-“Khoisan” peoples (Cashdan 1986; Barnard 1992; Blench 2006). Intriguingly, the possibility of a historical link to the Bantu has only rarely been considered (Westphal 1963; Cashdan 1986).

Located at the southwestern edge of the Bantu expansion and at the northwestern fringe of an area traditionally inhabited by Kx’a-speaking hunter-gatherers, the Angolan Namib desert forms a contact zone that mirrors the high variability currently observed in the wider region of southern Africa (Fig. 1). The dominant populations are the Himba and Kuvale, two matrilineal pastoralist populations commonly considered to be part of the broad Herero ethno-linguistic division that arrived in the area during the Bantu expansions, but whose relationships to one another and to other southwestern African Bantu speakers are not clear (Westphal 1963; Gibson 1977; Coelho et al. 2009; Barbieri et al. 2014b). In the orbit of these two groups gravitate several small-scale communities, including the Tjimba, the Kwepe, the Kwisi and the Twa, who share physical similarities and a matriclanic social organization with their Bantu neighbors, but whose origins remain unknown. Due to their patron-client relationship with the Himba and Kuvale, they are perhaps best described as peripatetic peoples (Bollig 2004), a category that encompasses small-scale, low-status, endogamous communities that are primarily non-food producing and provide specialized goods and services (e.g., as blacksmiths, healers, sorcerers) to their dominant neighbors. However, previous hypotheses about the history of the area, based on anthropological and linguistic data, suggest that these peripatetic communities are associated with very different migratory movements. The Kwisi and the Twa, who speak the Bantu language Kuvale, claim to be the native peoples of the Angolan Namib and have been considered remnants of the same set of pre-Bantu foraging populations to which the Damara were also ascribed (Almeida 1965; Estermann 1976). Their original language would have been lost after contact with the Bantu, similar to what has been claimed for the Pygmies of West and Central Africa (Güldemann 2008; Bahuchet 2012). The Kwepe are small stock breeders who until recently spoke Kwadi, a language that has been replaced by Kuvale and is now virtually extinct (Westphal 1963; Almeida 1965). Their linguistic heritage led to the proposal that they might represent a remnant group from the hypothetical Khoe-Kwadi migration introducing pastoralism to southern Africa (Güldemann 2008). Finally, the Tjimba are often considered Himba who lost their cattle but retained their language and other aspects of their culture (Warmelo 1951). Still, it has been suggested that some isolated Tjimba communities from Namibia might be connected to a more ancient foraging tradition (MacCalman and Grobbelaar 1965).

**Fig. 1.**
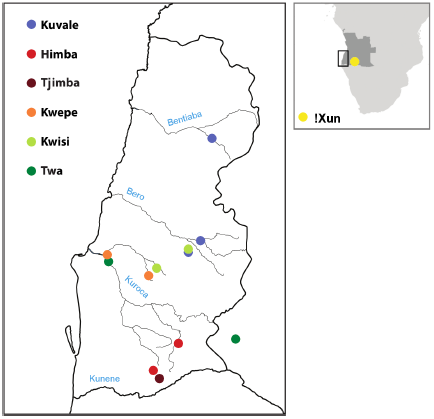
Map of sampling locations. Each location is colored by the corresponding population. On the right, Angola is highlighted in dark grey. On the left, an expanded view of the Angolan Namib (bold contour) is shown. The names of the main intermittent rivers are shown in blue.

All of these hypotheses entail a set of testable expectations about the genetic, linguistic and cultural relationships of the peoples living in the Angolan Namib. Specifically, from a genetic perspective it is expected that: i) the Kuvale and the Himba are related to each other as well as to other Herero-speaking peoples of southern Africa; ii) the Twa and the Kwisi are genetically similar to each other, but clearly distinct from their Bantu neighbors; iii) the Kwepe share genetic similarities with Khoe-speaking peoples from other regions of southern Africa; and iv) the Tjimba are either closely related to the Himba or have a very distinct genetic composition presumably related to the Twa and Kwisi. In this scenario, it is also expected that the matrilineal descent-group systems of the peripatetic peoples are relatively recent and were borrowed from their Himba and Kuvale neighbors, considering that matrilineality is known to be a distinctive feature of Bantu societies in southwestern Africa (Estermann 1952; Gibson 1956; Bollig 2006).

To date, the remote geographical location and high mobility status of the peripatetic peoples of the Angolan Namib have made it difficult to evaluate these predictions. Recently, in the course of field research being conducted in the area, we have located and contacted several communities belonging to the Twa, Kwisi, Kwepe and Tjimba ethnic groups who live in close proximity to the Kuvale and Himba populations. Here, we report for the first time a multidisciplinary assessment of the relationships between these populations based on newly collected linguistic data and 295 complete mtDNA genomes. Our results suggest that the maternal genetic structure of the different ethnic groups dwelling in the Namib Desert is largely derived from endogamous Bantu peoples and was strongly shaped by their matriclanic social organization, with contributions of non-Bantu populations being mostly restricted to “Khoisan” lineages. In this context, we propose that the social stratification and different subsistence patterns found in the area are not indicative of remnant groups, but reflect Bantu-internal variation and ethnogenesis.

## MATERIALS AND METHODS

### Samples

We analyzed 295 whole mitochondrial genomes from six populations living in the Namib desert (77 Himba; 85 Kuvale; 37 Kwepe, 24 Kwisi; 18 Twa; 15 Tjimba) and from 39 Kx’a-speaking !Xun hunter-gatherers from the Kunene Province (Fig. 1; Table S1). At all sampling locations, the purpose of the study was explained with the aid of bilingual native speakers. For each participant, we collected a saliva sample and information about language, matriclan and place of birth, up to the grandparental generation. With the exception of the !Xun, who do not have a clanic system, all sampled individuals identified as members of one out of 13 distinctive matriclans. Additional genealogical information, including relatedness with other donors, was also recorded. Given the intrinsic social structure of these highly endogamous groups, we only avoided including siblings and mother-offspring pairs in the final dataset (see Pinto et al. 2016 for details). The linguistic analyses were based on lexical data collected from individuals belonging to each sampled group, including two elder community members of the Kwepe community, who still remember Kwadi (see Pinto et al. 2016). As previously described (Pinto et al. 2016), the saliva samples, as well as the linguistic and the personal information, were collected with the donors' written informed consent in the framework of a collaboration between the Portuguese-Angolan TwinLab established between CIBIO/InBio and ISCED/Huíla Angola, with the ethical clearance of ISCED and the CIBIO/InBIO-University of Porto boards, and the support and permission of the Provincial Governments of Namibe and Kunene.

### mtDNA sequencing

Multiplexed sequencing libraries were produced from genomic DNA and enriched for mtDNA sequences following Meyer and Kircher (2010) and Maricic et al. (2010) with some modifications as detailed in Barbieri et al. (2012). The sequencing was performed on the Illumina Miseq platform with paired-end runs of 214 or 314 cycles. Base calling was performed with Bustard, adapters trimmed with leeHom (Renaud et al. 2014) and reads demultiplexed using deML (Renaud et al. 2015). The reads were aligned against the human reference genome 19 using a customized version of BWA v0.5.10-evan (https://bitbucket.org/ustenzel/network-aware-bwa; Li and Durbin 2009). Reads that aligned to the mitochondrial genome and known nuclear insertions of mitochondrial DNA (numts) (Li et al. 2012) were re-aligned to the mtDNA revised Cambridge Reference Sequence (Andrews et al. 1999) using BowTie2 (Langmead and Salzberg 2012), and the consensus sequences were called using an in-house script for detecting mtDNA heteroplasmies (Li and Stoneking 2012). The resulting mitochondrial genomes have a mean coverage of 400x. Missing nucleotides were replaced with the nucleotide that was present in all otherwise identical haplotypes of the dataset. With this imputation approach the missing data of the whole dataset (1057 missing nucleotides distributed across 10 samples) was reduced to 3 missing sites in a single sample. The Haplogrep webtool and Phylotree Build 16 were used to assign the haplogroup of each sample (Oven and Kayser 2008; Kloss-Brandstätter et al. 2011). Sequence alignments were performed with MUSCLE v.3.8 (Edgar 2004). The two poly-C regions (np 303-315, 16183-16194) were removed in all further analyses. Sequences are available from GenBank with accession numbers XXXXXXXX – XXXXXXXX.

### Genetic data analysis

Analyses of Molecular Variance (AMOVA), pairwise Φst values and genetic diversity indices were computed in Arlequin v3.5.2.2 (Excoffier and Lischer 2010). Non-metric multidimensional scaling (MDS) and k-means analyses based on pairwise Φst distance matrices were carried out in R, using the functions “isoMDS” from the package MASS (Venables and Ripley 2002) and “kmeans” with several random starts (Hartigan and Wong 1979), respectively. An additional matrix describing the relationships between populations solely on the basis of matriclan frequencies was generated in Arlequin v3.5.2.2 using a Fst-like distance treating different clans as alleles from a single locus. The correlation between genetic and clanic distances was assessed by performing a Mantel test with 1000 permutations of matrix elements to determine significance. Neighbor-Joining trees were generated using the R function “nj” from the package “ape” (Paradis et al. 2004).

Median-joining networks (Bandelt et al. 1999) were computed with Network 5.0 (www.fluxus-engineering.com) and customized in Network Publisher v2.1.1.2. The time to the most recent common ancestor (TMRCA) of subhaplogroups was estimated with Network from the *rho* statistic (Forster et al. 1996), using a mutation rate of 1.665 × 10 ^−8^ substitutions per nucleotide per year (Soares et al. 2009). The root defining the ancestral haplotype in each subhaplogroup was identified by using the full mtDNA network.

For population-based and sequence-based comparisons, we compiled a dataset comprising approximately 2,500 previously-published whole mitochondrial genomes from different regions of Africa (Table S2).

Probabilities of alternative evolutionary models were computed by using an Approximate Bayesian Computation (ABC) approach (Beaumont et al. 2002). For each model, two million datasets of complete mtDNA genomes were simulated assuming a mutation rate of 1.665 ×10 ^−8^substitutions/nucleotide/year (Soares et al. 2009), a transition bias matching the ratio observed in the empirical data, and a generation time of 28 years (Fenner 2005). Simulations were performed with fastsimcoal v2.5.2.1.1 (Excoffier et al. 2013) and summary statistics computed with Arlequin v3.5.2.2 (Excoffier and Lischer 2010), both within the framework of ABCtoolbox (Wegmann et al. 2010). The summary statistics used for comparing the observed and simulated data were the number of haplotypes (k), sequence diversity (H), number of segregating sites (S), number of private segregating sites (prS), Tajima's D (D) and mean number of pairwise differences (MPD), all computed within populations. In addition, population pairwise Φst and pairwise MPD was computed between pairs of populations. All summary statistics were standardized.

The ABC estimations were performed with a general linear model (GLM) regression adjustment (Leuenberger and Wegmann 2010; Wegmann et al. 2010) applied to the 10,000 retained simulations (0.5%) closest to the observed data. Model selection was based on posterior probabilities estimated using the marginal density of each model relative to the density of all models. The power to correctly select a given model was assessed by using 1,000 pseudo-observed datasets taken from that model and calculating the number of times it had the highest posterior probability when compared with alternative models (Veeramah et al. 2012). To reduce the effects of including summary statistics that are redundant or do not capture the main features of the data, we additionally performed model selection using a subset of summary statistics that were only moderately correlated (Pearson's r ^2^ < 0.8) and exhibited the highest power to discriminate between models, as proposed by de Filippo et al. (2016) (Table S3).

To estimate parameters from the most supported model, we transformed summary statistics from simulated and observed data into partial least squares (PLS) using the R scripts provided in ABCtoolbox (Wegmann et al. 2009, 2010). The smallest set of PLS components with the largest amount of information about the model parameters was selected by using Root Mean Square Error (RMSE) plots (Wegmann et al. 2009). The estimation was then performed as described above and the posterior distributions of individual parameters were checked for bias. We randomly selected 1,000 pseudo-observed datasets generated with known parameter values to determine the coverage of the posteriors (the proportion of times a true parameter value is present in a given credible interval) (Wegmann et al. 2009; Wegmann and Excoffier 2010). A Kolmogorov-Smirnov test was applied (with Bonferroni correction) to assess the uniformity of the posterior quantiles. To determine the power of parameter estimation, we computed the coefficient of variation R ^2^ by regressing the PLS components against each model parameter (Neuenschwander et al. 2008). To evaluate the accuracy of the mode as a point estimate, we calculated the RMSEmode for each parameter based on 1,000 pseudo-observed datasets (Wegmann and Excoffier 2010). Pairs of PLS components from the retained simulations were plotted together with the transformed observed data in order to check how well the retained simulations fit the observed data.

### Linguistic data analysis

We collected data from southwestern Bantu languages of southwestern Angola, as well as comparative samples from Kwadi (as remembered by two Kwepe elders) and the !Xun variety of the Kunene Province. The linguistic data from Bantu are based on a 600-item wordlist that is a subset of the Summer Institute of Linguistics (SIL) Comparative African Wordlist (Snider and Roberts 2006) and was supplemented by comparative material from Namibian Herero (Möhlig and Kavari 2008), and several varieties belonging to the Nyaneka-Nkhumbi cluster (Humbe, Muhila, Nyaneka, Ngambwe, Handa) (unpublished data from Jordan and Manuel I 2013; Jordan 2015).

Following an analysis of regular sound correspondences, we established 693 cognate sets based on 273 meanings, which include the Swadesh 200 (Swadesh 1952) and Leipzig-Jakarta wordlists (Haspelmath and Tadmore 2009), minus function words, personal pronouns, and question words. For computational purposes, we coded languages for presence (1) or absence (0) of a particular lexical root. As our data from Himba and Tjimba displayed a high degree of linguistic homogeneity, they were combined and treated under the label “Himba”. Based on our coded dataset, we generated a matrix of linguistic distances (1 minus the percentage of cognate sharing) and computed a Neighbor-Joining tree using the R package “ape”, as described above. Linguistic distances were compared with genetic distances with a Mantel test, as described above.

We further used a Bayesian phylogenetic approach as implemented in the BEAST2 framework (Bouckaert et al. 2014) and tested three models included in the Babel package (Bouckaert R 2016): (1) Continuous Time Markov Chain (CTMC); (cf. Greenhill and Gray, 2009); (2) Covarion (Penny et al. 2001; Atkinson et al. 2008); (3) Dollo (Nicholls and Gray 2006). We ran an analysis for each model, with a chain length of 10,000,000, sampling every 1000 steps. The first 100,000 steps were discarded as burn-in.

To evaluate the performance of these models with our dataset, we used the Tracer software (Rambaut and Drummond 2007) to compare the Akaike Information Criteria through Markov chain Monte Carlo (AICM) of each analysis, where lower AICM values indicate better model fit (Baele et al. 2013). We found that the model displaying the best fit for our data was Covarion (AICM = 7116), outranking both CTMC (AICM = 7146) and Dollo (AICM = 7682).

The output of the analysis was visualized in DensiTree (Bouckaert 2010) in order to display reticulations and conflicting signals.

## RESULTS

### Genetic and matriclanic diversity in the Angolan Namib

By performing an analysis of molecular variance, we found that 25.2% of the total genetic variation in our sample is due to differences between populations. This level of genetic differentiation is 20.2% even when the ! Xun are removed and is higher than previously observed among Bantu (5.5%; Barbieri, et al. 2014b) and “Khoisan” populations (16.6%; Barbieri, et al. 2014a). The levels of intra-population diversity are highest in the Kuvale and Himba (mean value of haplotype diversity, 0.95) and lowest in in the Kwepe (0.67), who display only five different haplotypes (Table S1).

A non-metric multidimensional scaling plot (MDS) based on pairwise Φst distances reveals three main vertices of divergence (Fig. 2a): i) the !Xun from Kunene Province, who have high frequencies (97%; Table S4) of haplogroups L0d and L0k that typically predominate in most “Khoisan” populations from southern Africa (Barbieri et al. 2014a); ii) the Tjimba and Himba, whose close genetic relationship supports the view that the two groups are merely distinguished by their socio-economic status (Warmelo 1951; Vashro and Cashdan 2015); iii) the Kwisi and Twa, whose genetic proximity is consistent with previous claims that these communities represent northern and southern branches of the same ethnic group respectively (Estermann 1976). The differences in the mtDNA composition of the Namib peoples are mainly due to the uneven distribution of nine common subhaplogroups that collectively account for 90% of their observed variation, each with a small number of haplotypes rarely exhibiting more than 5 pairwise differences (Figs. 2b and 3; Table S4): L0a1b1, L0a1b2, L0a2a1b and L1c1b are very common in the Kwisi and Twa; L3e1a2 and L3d3a1a predominate in the Himba and Tjimba, while L0d1a1b1a and L0d1b1b are very frequent in the Kuvale, placing them closer to the ! Xun (Fig. 2a). With the exception of the Kwisi, L3f1b4a is found at relatively high frequencies in most groups. An assessment of lineage sharing among different populations shows that the most common subhaplogroups among the Himba/Tjimba and Kuvale (Fig. S1) are rarely found in other groups, except for one single L3f1b4a haplotype that is very frequent in the Kwepe but is likely to have originated in the Himba, who display a higher L3f1b4a diversity (Fig. S1). Conversely, haplotypes belonging to subhaplogroups that are frequent and diverse in the Kwisi, the Twa or the Kwepe can be found at moderate frequencies in the Himba and Kuvale (Fig. S1), suggesting that gene flow occurs preferentially from these peripatetic communities into the dominant groups.

**Fig. 2.**
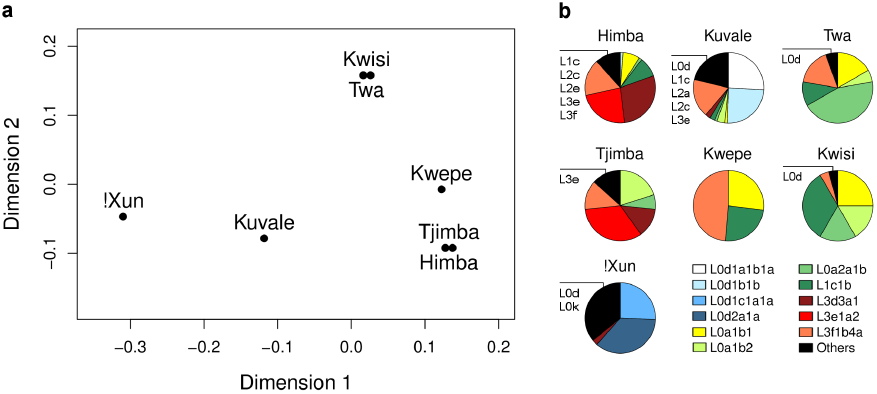
Multidimensional scaling analysis and haplogroup variation in southwestern Angola. (a) MDS plot based on Φst genetic distances. The pairs Kwisi-Twa and Tjimba-Himba are not significantly different, with *p*-values 0.11 and 0.16, respectively. Stress value: 0.006. (b) Frequencies of the most common subhaplogroups (≥ 20% in at least one population) are shown for each population. The remaining subhaplogroups are pooled under the category "Others" (black), with the major haplogroup assignments within this category listed for each population. Note that major haplogroups that are represented in the plots by a specific subhaplogroup, might appear again in the category "Others’ to indicate other low frequency subhaplogroups.

The nine most common subhaplogroups are associated with all 13 matriclans identified during our survey, with the number of clans in each subhaplogroup varying from one to five (Fig. 3). The occurrence of several clans in the same subhaplogroup has several potential explanations, including adoption, patrilineal transmission, or chance. However, this pattern can also be explained by a well documented Herero custom of splitting the same line of descent into different clans, forming clan-groups with a claimed common ancestor designated as phratries (Gibson 1956; Vivelo 1977). Interestingly, we found that three pairs of clans that were reported to us has sharing a distant ancestor were also associated with the same subhaplogroup: Mukwalukune / Mukwatjiti (L0d1a1b1a); Mukwanambula / Mukwangombe (L0d1b1b and also L0a2a1b) and Mukwandjata / Mukwambua (L3f1b4a).

**Fig. 3.**
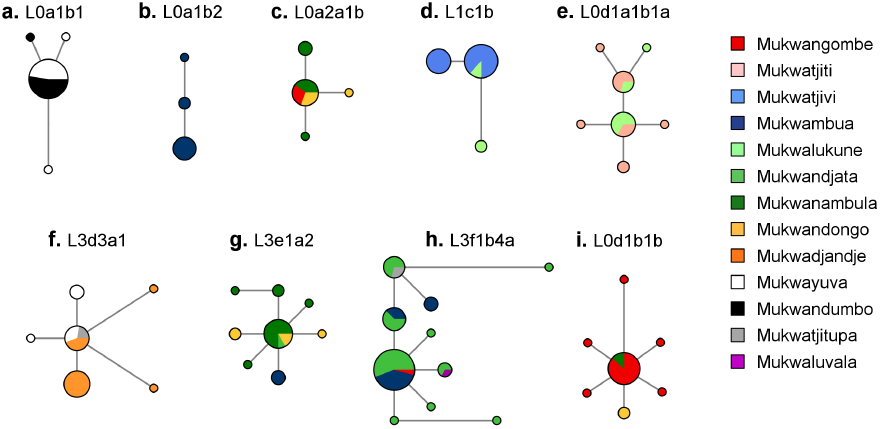
Median-joining networks showing haplotype variation within the most common subhaplogroups of the Angolan Namib. Circles represent mtDNA haplotypes, with size proportional to frequency and color corresponding to clan affiliation. Line lengths are proportional to the number of mutational steps. Indels were not included.

While most clans are distributed across multiple populations (Fig. 4a), we found several cases where the same clan is associated with different subhaplogroups in different populations (Fig. S2b, e, g, h, k), suggesting that clan sharing is not always due to migration. All these cases involve at least one common subhaplogroup from a dominant population (Kuvale or Himba), and one common subhaplogroup from the Twa, Kwisi and Kwepe peripatetics.

**Fig. 4.**
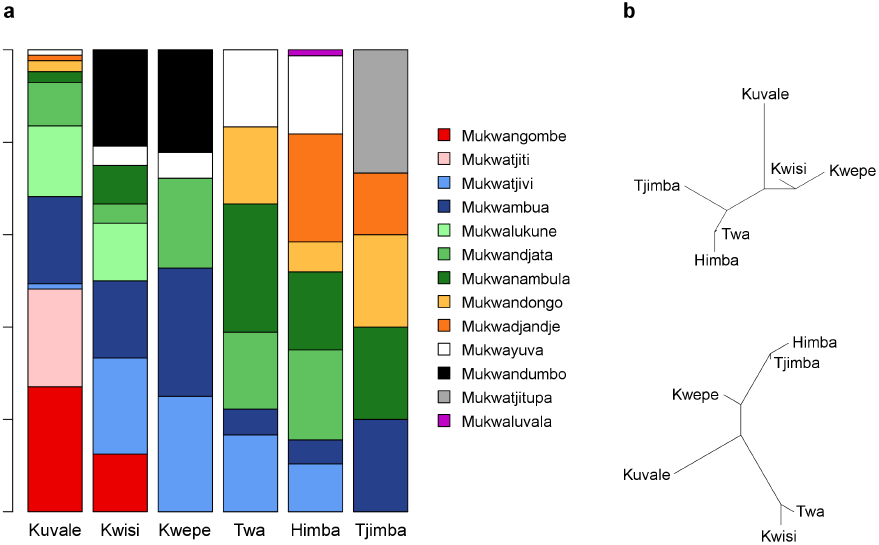
Relationship between genetic and clanic distances in populations of the Angolan Namib. (a) Matriclan distribution within each population. (b) Neighbor-joining tree based on clan distances (top) and Φst genetic distances (bottom).

The absence of a one-to-one correspondence between matriclans and subhaplogroups decreases the association between the distribution of matriclans and the genetic differentiation among populations (Fig. 4b): in some cases, subhaplogroups that are associated with the same matriclan predominate in populations that are genetically very divergent; in others, subhaplogroups that are shared across genetically similar populations are associated with different matriclans. Consequently, distance matrices between populations based on matriclans and mtDNA are clearly uncorrelated (Mantel test *p* = 0.57; Fig. 4b).

In spite of these exceptions, we found that as much as 51% of the total mtDNA variation reflects differences between matriclans, a highly significant value (AMOVA; *p* < 0.00001) that is more than two times greater than the 20.2% proportion calculated among ethnic groups, indicating that there are remarkable differences in the mtDNA sequence profiles of individual matriclans (Table S5).

Moreover, as shown in Figure 5a, the distributions of pairwise differences clearly indicate that mtDNA sequences drawn from the same clan have a significantly higher average probability of being closely related (<= 5 pairwise differences) than two sequences randomly sampled from the whole Namib pool (0.53 vs. 0.10; *p* < 0.001, Fisher exact test), or from the same population (0.53 vs. 0.18; *p* < 0.001), indicating that individuals from the same clan are more likely to share a subhaplogroup. This association becomes even stronger when mtDNA sequences are sampled in the same clan and the same population (0.53 vs. 0.63; *p* < 0.001).

**Fig. 5.**
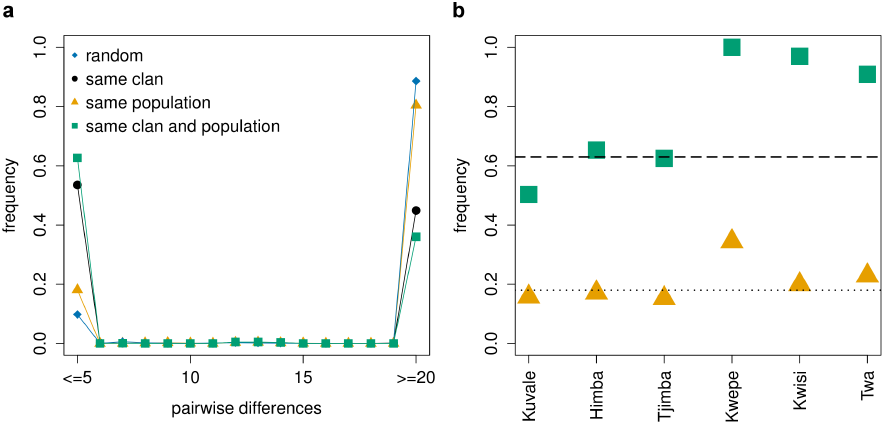
Genealogical consistency of matriclans. (a) Distribution of pairwise differences obtained by randomly drawing pairs of sequences from: i) the whole Angolan Namib pool, ii) the same population, iii) the same clan, and iv) the same clan and population. (b) Sequence similarity in Angolan Namib populations computed for pairs of sequences randomly drawn from each population (orange triangles) and from individuals belonging to the same clan in each population (green squares). The dotted and dashed lines show the average sequence similarity computed within populations, regardless of the clan, and within clans, respectively. Sequence similarity was measured by the frequency of sequence pairs with ≤ 5 differences.

The probability of sampling related sequences within clans is significantly elevated in all populations (Fig. 5b; *p*< 0.001 in all comparisons), and is remarkably high in the Kwepe, Twa and Kwisi, who display greater levels of within-clan sequence similarity than the Tjimba, Himba and Kuvale.

To rule out the possibility that close kin relationships could have inflated the likelihood that individuals from the same clan have exactly the same haplotype, we restricted the analysis to closely-related but non-identical haplotypes. We randomized one million times the matriclan labels on observed matriclan/haplotype pairs and then calculated the probability of finding within the same matriclan two haplotypes with 1 to 5 differences (Fig. S3). As shown in Table S6, in most groups the observed value for this probability is too high to be obtained by chance, indicating that similar (but non-identical) haplotypes have a high probability of sharing clans by inheritance. The only non-significant values were found among the Kwepe and the Tjimba, whose low levels of within haplogroup diversity reduce the power of the test (Fig. 2b; Fig. S1). Note that by using this approach we made the conservative assumption that all individuals within the same matriclan/haplotype pair share a common ancestor, which drastically reduces the number of independent matriclan assignments that are needed to randomly match the observed data (Fig. S3).

Table S7 presents the estimates of the times to the most recent common ancestors (TMRCA) of the nine predominant subhaplogroups obtained with the *rho* statistic (Forster et al. 1996). Due to the association between clans and subhaplogroups exhibited by most populations, these TMRCAs can be used as proxies for the coalescent ages of the oldest clans in each clan-group. However, it is not possible to provide separate estimates for matriclans associated with the same subhaplogroup, since these clans often share the TMRCA of the whole subhaplogroup and represent different samples from the same genealogy (Fig. 3). The TMRCA estimates range from ~560 to ~3,140 years (average ~1800 years) with large standard deviations.

### Relationships with other populations

When the genetic profiles of the populations from Namib are compared with an extended mitochondrial genome-dataset including other groups from Angola (Nyaneka-Nkhumbi, Ovimbundu, Ganguela) and the wider region of southern Africa (Fig. 6), the Kwisi and the Twa remain outliers, while the Tjimba and Himba fall close to the Herero, Himba and Damara from Namibia (see also Soodyall and Jenkins 1993; Barbieri et al. 2014a; Barbieri et al. 2014b). The Kuvale, in contrast, are more similar to other populations with high levels of maternal Bantu-“Khoisan” admixture, including the Tshua, Shua, TcireTcire and ∥Ani. The Kwepe are not close to any Khoe-speaking group, even though they spoke the related Kwadi language until recently (Almeida 1965; Pinto et al. 2016). The !Xun from the Angolan Kunene Province are related to Kx’a- and Tuu-speaking groups from Namibia and Botswana.

**Fig. 6.**
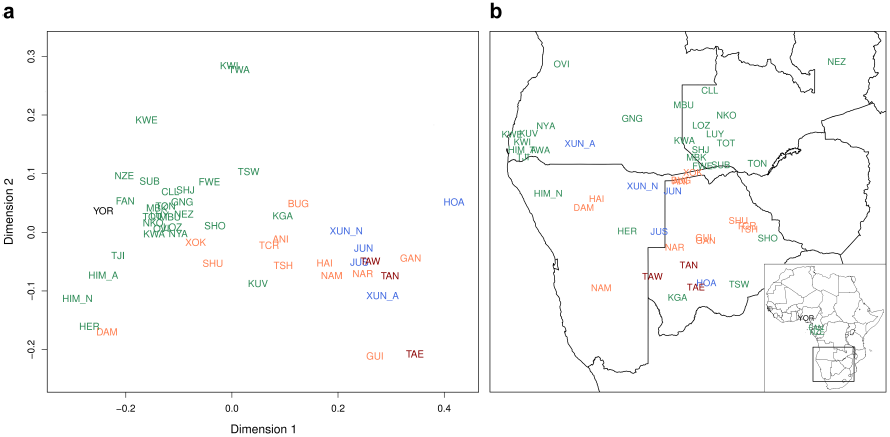
Multidimensional scaling analysis in the wider region of southern Africa. Colors correspond to language families: Niger-Congo non-Bantu (black), Niger-Congo Bantu (green), Kx’a (blue), Tuu (dark red), KhoeKwadi (orange). The code used for each population can be found in Table S2. (a) MDS plot based on Φst genetic distances. Stress value: 9.4.(b) Geographic origin of the populations included in the MDS analysis.

These patterns are confirmed and complemented by the clustering results obtained with the k-means algorithm (Fig. S4). With the exception of the Kuvale, all the populations from Namib are initially lumped into a cluster encompassing most Bantu-speaking peoples (k=2 in red). Further partitions: i) isolate a homogeneous group of Bantu-speaking populations that forms a central core in the MDS plot (k=4 in green); ii) separate the Twa and Kwisi from the other clusters (k=6 in yellow); and iii) group the Angolan Himba with the Herero, Himba and Damara from Namibia (k=7 in orange). An outstanding feature of the k-means partitions is the wide dispersal across different clusters of the Khoe-Kwadi-speaking populations represented in our dataset. Some groups from the Central Kalahari (Gui, G?ana and Naro) and Namibia (Nama and Hai?om) cluster together with Kx’a and Tuu-speaking “Khoisan” peoples (k=2-7). Groups from the eastern Kalahari (Tshwa, TcireTcire) and Okavango (?Ani and Buga) form a cluster with high levels of maternal Bantu/“Khoisan” admixture together with the Bantu-speaking Kuvale, Tswana and Kgalagadi (k=3-7). Finally, the Damara, the ?Xokhoe and the Kwepe (formerly speaking Kwadi), in spite of their high levels of genetic differentiation, are grouped together with Bantu-speaking populations that have low amounts of “Khoisan” admixture (k=2-7).

The phylogeographic analysis of the mtDNA lineages from the Namib populations provides additional information about their relationships with groups from adjacent areas (Fig. S5). Subhaplogroups L1c1b and L0a1b2, are remarkable for their molecular divergence and geographical confinement to southwestern Angola (Fig. S5c, i). Other major subhaplogroups have molecularly close neighbors in several Bantu-speaking populations from southern Africa (L0a2a1b and L3f1b4a; Fig. S5d, l) or are related to sequences that are mostly shared by Bantu and Khoe-Kwadi groups from the area (L0a1b1, L3d3a1 and L3e1a2; Fig. S5b, j, k). None of the L0d lineages common in the Kuvale (L0d1a1b1a and L0d1b1b) were found in the !Xun from Angola, the nearest “Khoisan” group from the Namib desert. Instead, their L0d1a1b1a haplotypes, also observed in the Himba from Namibia, are close to lineages that were found in the Khoe-Kwadi-speaking Shua from Botswana, while the L0d1b1b haplotypes are remotely related to sequences observed in the Damara from Namibia and the Luyana from Zambia (Fig. S5e, f). The most common subhaplogroups in the !Xun (L0d1c1a1a, 26% and L0d2a1a, 36%) have unique haplotype matches with !Xun and Ju|'hoan from northern Namibia and display sequences that are closely related to “Khoisan” groups from southern Africa (Fig. S5g, h). Taken together, these results indicate that, with two exceptions (L1c1b and L0a1b2), most sequences from southwestern Angola are nested in the phylogeographic pattern that emerged from the contact of previously identified population strata from southern Africa.

### Testing relationships of Kuvale and Herero/Himba/Damara

As previously noted (Barbieri et al. 2014b), the close proximity of the Himba and Herero pastoralists to the Damara, who speak the same Khoe language as the Nama and have a peripatetic lifestyle, stands in stark contrast to their genetic distinctiveness from the linguistically and culturally similar Kuvale. Based on resampling tests, Barbieri et al. (2014) suggested that the sharing of a common ancestry by the Herero, Himba and Kuvale was not compatible with a scenario of shared ancestry between the Herero, Himba and Damara. Here, we address this question by lumping the closely related Herero, Himba and Damara (all clustered by k- means at k=7; Fig. S4) into a single metapopulation (HHD) and testing three evolutionary scenarios relating this metapopulation with the Kuvale and two neighboring populations (Nyaneka-Nkhumbi and !Xun), using an Approximate Bayesian Computation (ABC) approach (Beaumont et al. 2002). The !Xun-speaking “Khoisan” from Angola were always used as an outgroup and we assumed that their split predated all other events (Fig. 7). The Nyaneka-Nkhumbi provide a southwestern Bantu-speaking reference population located to the northeast of the Namib desert (Fig. 6b). In the first scenario, an early divergence of the Kuvale is followed by a more recent split between the Nyaneka-Nkhumbi and the HHD metapopulation (Fig. 7, Model A). The second scenario postulates a recent common origin of the Kuvale and HHD (Fig. 7, Model B). The third scenario assumes that the most recent common origin is between the Kuvale and the Nyaneka-Nkhumbi (Fig. 7, Model C). Asymmetric migration was allowed between all pairs of populations. Priors for splitting times (T), effective population sizes (Ne) and migration rates (m) are shown in Table S8. The power to predict the correct model was 0.47, 0.48 and 0.44, in simulated models A, B, and C, respectively. These values are significantly different from the expected 0.33 if there was no discriminatory power (*p* < 0.001, binomial test).

**Fig. 7.**
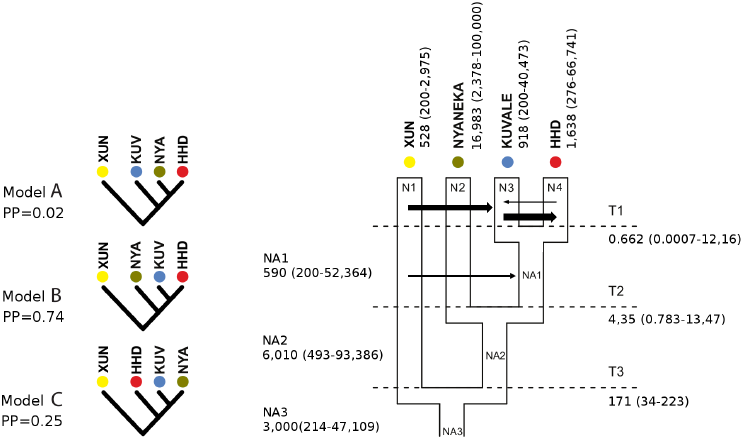
Demographic models tested by ABC. The three tested models are shown on the left with their respective posterior probabilities (PP). Migration ratios above 0.0001 or effective migration (Nm) above 2 are represented in the plot by arrows with width proportional to Nm. NA1- NA3: ancestral effective population sizes; N1 - N4: current effective population sizes; T1-T3: divergence times.

Model B, assuming a recent common origin of the Kuvale and HHD, was the most supported scenario, with a posterior probability of 0.74 (Fig. 7, Model B). By iteratively excluding summary statistics that were highly correlated (Pearson's r ^2^ > 0.8), starting with those which had less power to discriminate between models (de Filippo et al. 2016), we found that model B was still the most supported model.

We additionally used the ABC framework to estimate the demographic parameters of the best supported scenario based on 2 million simulations (Fig. 7, Model B; Table S8; Fig. S6). Assuming a generation time of 28 years (Fenner 2005), our estimate for the time of split of the !Xun (T3; ~170 kya; 95% CI: 34-223 kya) is consistent with previous calculations of the divergence time of “Khoisan” peoples from other sub-Saharan African populations (Behar et al. 2008; Schlebusch et al. 2012; Veeramah et al. 2012). The proposed time of split between the Kuvale and HHD (T1) is quite recent (0.662 kya; 95% CI: 0.001-12.16 kya), while the date of divergence of the Nyaneka-Nkhumbi (T2; 4.35 kya; 95% CI: 0.783-13.47 kya) is probably overestimated, given the available archeological evidence for the arrival of Bantu peoples in southern Africa of only about 1.5 kya (Russell et al. 2014).

Our estimates of Ne show that the Nyaneka-Nkhumbi have the largest effective population size (~17,000; 95% CI: 2,378-100,000), followed by the HHD, the Kuvale and the !Xun, with estimates of ~1,600 (95% CI: 276 −66,741), ~900 (95% CI: 200-40,073) and ~500, respectively (95% CI: 200-2,975) (Fig. 7; Table S8). The point estimates of ancestral effective population sizes (NA1 and NA2) suggest that the Nyaneka-Nkhumbi experienced a ~3-fold growth after their split (Ne A2 = 6,000 to Ne Nyaneka-Nkhumbi = 17,000), while the size of the ancestors of the Kuvale and HHD underwent a ~10-fold reduction (Ne A2=6,000 to Ne A1 = 600) (Fig. 7; Table S8).

The migration estimates, expressed either as the proportion of immigrants in a population per generation (m) or the absolute number of immigrants per generation (Nm), show that the amount of gene flow into the !Xun is negligible (Table S8), in agreement with their genetic proximity to other “Khoisan” groups and their pronounced divergence from the HHD and Kuvale (Fig. 6a). Elevated migration rates from the !Xun into the Kuvale (m = 0.021; Nm = 18.9 migrants/generation), are in accordance with the high amount of characteristic L0d haplotypes that was found in this population (Figs. 2 and 7; Table S8). However, this result should be interpreted with caution since most L0d lineages in the Kuvale belong to two subhaplogroups that are likely to be derived from only two ancestral women (Table S4; Fig. S1), and probably were not transferred by the continuous gene flow process simulated in our ABC analysis. We additionally estimated high migration rates from the !Xun to the common ancestor of the Kuvale and HHD (m = 0.010, Nm = 5.7), from the Kuvale to HHD (m = 0.015; Nm = 24.5), and to a lesser extent from the HHD to the Kuvale (m = 0.005, Nm = 4.2) (Fig. 7).

### Linguistic analyses

The high amount of genetic divergence among the Namib peoples (Fig. 2a) contrasts with the relative linguistic homogeneity of the area, where all groups presently speak either Himba or Kuvale. While the classification of Himba as a variety of the Herero language is fairly straightforward and widely accepted, the position of Kuvale is less clear (Westphal 1963; Vansina 2004; Maho 2009). Moreover, the Bantu languages spoken by the Kwisi and Twa have long been the subject of speculation (Westphal 1963). To evaluate the relationships between the Himba and Kuvale languages that are currently spoken in the Angolan Namib, as well as their links to Namibian Herero and to Nyaneka-Nkhumbi southwestern Bantu varieties, we first undertook a lexicostatistical analysis and calculated a language distance matrix based on 693 cognate sets. In the Neighbor-Joining tree based on the language distance matrix, Kuvale forms its own cluster, separated from Himba and Herero on one side and various dialects of Nyaneka-Nkhumbi on the other (Fig. 8a). Kuvale as spoken amongst the Kwisi and Kwepe is fully within the range of the cluster. The variety spoken by the Twa seems to have been influenced by Himba and lies in-between the Kuvale and the Himba/Herero clusters. Furthermore, based on a careful comparison of our Bantu wordlists with lexical data from Kwadi (Westphal 1963, supplemented by our own field notes), Khoe (Vossen 1997) and !Xun (König and Heine 2008) we note that no linguistic variety spoken in Namib displays any lexical peculiarities that could be linked to influence from a non-Bantu substrate. As a result of the nesting of linguistic varieties spoken by the peripatetic Kwisi, Twa and Kwepe within the range of Kuvale and Herero/Himba, distance matrices based on linguistic and genetic distances between the Namib groups are uncorrelated (Mantel test, *p* = 0.18).

**Fig. 8.**
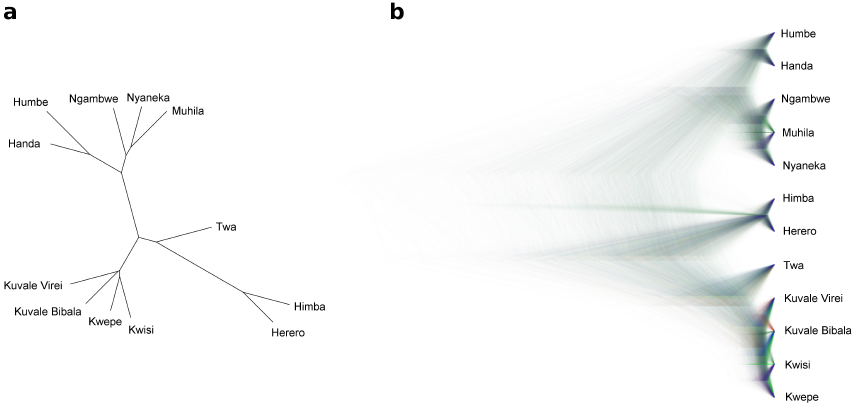
Linguistic relationships between Kuvale, Himba, Herero and Nyaneka-Nkhumbi. The Kuvale sample includes varieties spoken by the Kuvale people (Kuvale Virei and Kuvale Bibala), as well as the Kwepe, the Kwisi and the Twa. The Nyaneka-Nkhumbi sample includes varieties spoken by the Handa, Humbe, Ngambwe, Nyaneka and Muhila peoples. a) Neighbor-joining tree. b) Bayesian trees plotted with DensiTree.

To gain a better understanding of the historical relations between Nyaneka-Nkhumbi, Herero and Kuvale, we additionally undertook a Bayesian phylogenetic analysis in BEAST, using the same 693 cognate sets underlying the Neighbor-Joining tree in Figure 8a. As in our previous analysis, all language clusters (Herero, Kuvale and Nyaneka-Nkhumbi) are unequivocally identified (*p* = 1.00; Fig. 8b; Fig. S7). The analysis further suggests a more recent common ancestor for Herero and Kuvale (*p* = 0.9) than either language shares with the five varieties of Nyaneka-Nkhumbi we included in our analysis. This result is remarkably congruent with Model B of the ABC analysis, which suggests that Kuvale and Herero are more closely related than either population is to Nyaneka-Nkhumbi (Figs. 7 and 8b). Within Kuvale, we find no well-supported subclusters, except for the initial split from Twa (*p* = 1.00), which is grouped with the other varieties, but remains an outlier (Fig. 8a; Fig. S7).

## DISCUSSION

In recent years, a growing number of studies on the population history of southern Africa has considerably broadened our knowledge concerning the historical interactions of groups dwelling in and around the Kalahari Basin (Schlebusch et al. 2012; Pickrell et al. 2012; Barbieri et al. 2014b; Marks et al. 2015). Within this geographical area, the focus has largely been on “Khoisan”-speakers and the southeastern Bantu populations whose genetic and cultural make-ups are thought to have been shaped by contact with indigenous foragers and herders. In the Southwest, new genetic data have recently become available for populations from Namibia and southern Africa (Uren et al. 2016; Montinaro et al. 2017), while the groups to their north remain the subject of intense speculation, but constitute a noticeable gap in the available literature. Our study presents for the first time full maternal genomes and linguistic data from Angolan populations previously deemed inaccessible or vanished (Almeida 1965; Estermann 1976), including Bantu-speaking groups, as well as the formerly Kwadi-speaking Kwepe. We sampled both foraging and pastoral populations, placing special emphasis on the analysis of the coherence of the matriclanic system that characterizes the area and unites populations of different social status and modes of subsistence. In this framework, we are now able to address different historical hypotheses about the present-day diversity found in the Namib Desert both from a local perspective and within the context of the wider region of southern Africa.

### Genealogical consistency of matriclans

A remarkable feature of the social organization of all the populations from the Angolan Namib and other southwestern Bantu peoples is their matrilineal descent-group system in which individuals are affiliated to the clan of their mother, and members of the same matriclan (sing. *eanda*) consider themselves as distant relatives that descend from an unknown founder woman (Estermann 1952; Gibson 1956; Bollig 2006). Although some populations may have dual descent systems and additionally form patriclans, it is the matrilineal principle that regulates key aspects of community life, such as cattle inheritance, social obligations, marriage preferences and group membership (Gibson 1956). However, the consistency of southwestern African matriclans has been difficult to validate with genealogical data, since the relationships between members of the same clan are often considered to be too distant to be traced accurately (Gibson 1956; Vivelo 1977). Furthermore, it has been suggested that members of low-status peripatetic communities borrowed the matriclanic system from their dominant neighbors as a means to achieve better integration into the regional network of the southwestern Bantu societies (Estermann 1976; Bollig 2004).

In this study we relied on the maternal inheritance of mtDNA to show for the first time that matriclans are indeed good descriptors of deep genealogical relationships in pastoral and peripatetic Bantu-speakers from southwestern Angola. Several interrelated lines of evidence support this conclusion: i) a high proportion of the total mtDNA variation is found among matriclans (Φst = 0.51; *p* < 0.00001); ii) individuals from the same clan have a significantly increased probability of having related mtDNA haplotypes that are likely to belong to the same subhaplogroup (Figs. 3 and 5); iii) the average TMRCAs of major subhaplogroups (~1,800 years) suggests that the oldest matriclans are not recent and probably date back to the arrival of Bantu-speaking peoples to southern Africa.

In spite of this evidence, we found that several matriclans likely became associated to more than one subhaplogroup through multiple founders in different populations. Since these cases often involve a subhaplogroup restricted to the Himba or Kuvale and a subhaplogroup predominant in the Kwepe, Twa or Kwisi (Fig. S2b, e, g, h, k), it may be argued that these low-status peripatetic communities were clanless and recently borrowed the matriclanic system from their dominant neighbors, as proposed previously (Estermann 1976; Bollig 2004). However, such an imitation scenario is difficult to reconcile with the antiquity and the genealogical consistency of the matriclanic system observed in all peripatetic populations (Table S7; Fig. 5b). Furthermore, our permutation tests indicate that random assignment of clans, as would be expected in a borrowing situation, is very unlikely in these communities (Table S6).

Alternatively, we find it more plausible that the Twa, Kwisi and Kwepe may have had their own matriclan systems, and merely replaced their pre-existing clan labels with those of their dominant neighbors. This seems to be particularly evident among the Twa, who have a genealogically consistent matriclan system based on a clan inventory similar to the Himba, despite their close genetic relationship with the Kwisi (Fig. 4b). The cultural approximation of the Twa to the Himba, which might be driven by geographical proximity (Fig. 1), is also reflected in the apparent influence of Himba on the linguistic variety spoken by the Twa (Fig. 8), as well as the documented tendency of the Twa to mimic the distinctive attire of the Himba women (Estermann 1952). More generally, it is likely that clan-switching has facilitated female gene flow from the peripatetics into the dominant communities (Fig. S1), thus explaining the reduced levels of sequence similarity observed within Himba and Kuvale clans (Fig. 5b; Fig. S2; Table S6).

The matriclanic organization of the Namib peoples seems to have had a strong impact on their current patterns of mtDNA variation. The fact that the percentage of the total genetic diversity that is found between clans (Φst = 0.51) is much higher than that observed between populations (Φst = 0.20) suggests that ethnic groups arose from the assemblage of genetically different clans instead of clans being formed just by fissions occurring within groups. Thus, although both clan and group membership are determined by the mother, it is clear that the matrilineal principle is frequently violated during ethnogenesis. This pattern is especially striking among the Kuvale, who are highly endogamous and ethnically Bantu, yet comprise among their founders two descent groups (L0d1a1b1a and L0d1b1b; Figs. 2a and 3), including the powerful clan of the cattle (Mukwangombe), that ultimately trace their origin to “Khoisan” populations. This type of population structure closely mirrors patterns of Y-chromosome variation previously reported in traditional patrilineal societies from other regions of the world (Chaix et al. 2004, 2007; Sanchez-Faddeev et al. 2013).

### The southwestern African pastoral scene: Herero, Himba, Damara and Kuvale

The Himba and Kuvale from Angola are generally considered to be part of a broad cultural cluster of Bantu-speaking cattle herders that includes adjacent Himba groups from Namibia, as well as Herero populations extending from Namibia to Botswana (Bollig and Gewald 2009). Besides sharing many aspects of their pastoral culture, these peoples are commonly thought to speak dialects of the same Herero language, which has been grouped with Nyaneka-Nkhumbi and Ovambo into a division of southwestern Bantu referred to as Cimbabesia (Vansina 2004). However, the internal relations and migration routes of the southwestern Bantu herders, as well as the origins of their pastoral tradition remain poorly understood (Gibson 1977; Bollig and Gewald 2009).

Our results, together with previous work, show that the Himba, Tjimba and Herero share a mtDNA profile that sets them apart from the Kuvale and other Bantu-speaking populations, but is not significantly different from the Damara who speak the same Khoe-Kwadi language as the pastoral Nama (Figs. 2b and 6a) (Coelho et al. 2009; Barbieri et al. 2014b).

The most striking aspect of the Kuvale’s maternal heritage is the high frequency (~50%) of characteristic “Khoisan” lineages associated with sequence types (L0d1a1b1a and L0d1b1b) that are likely to be derived from only two ancestral women (Figs. 2a and 3). In contrast, the Himba, Herero and Damara have much lower frequencies of “Khoisan” mtDNA (10-17%), and share unusually high frequencies of subhaplogroup L3d3a (38- 61%), which is present in several Bantu, Kx’a and Khoe-Kwadi speaking populations of southwestern Africa (Fig. S5j; Soodyall and Jenkins 1993; Barbieri et al. 2014b).

Previous interpretations of this mtDNA pattern have proposed that L3d3a was a pre-Bantu lineage retained by the Damara that was subsequently transferred to the Himba and the Herero through admixture, instead of being inherited from a common ancestor by all three populations (Barbieri et al. 2014b). By using ABC analysis to explicitly test alternative evolutionary hypotheses about the relationships between the Kuvale, the Nyaneka- Nkhumbi and a meta-group lumping the Himba, Herero and Damara (HHD), we found that the maternal heritage of the latter group is nested within the southwestern Bantu peoples and shares a recent common ancestor with the Kuvale (Fig. 7). In this context, it seems likely that the HHD and Kuvale represent the southern and northern branches, respectively, of a proto-population whose origins may be tentatively placed to the east of their present locations on the basis of the geographic distribution of their most common DNA lineages (Fig. S5e, f, j, k). The separation between the HHD and the Kuvale is paralleled by our linguistic results, which show that the Kuvale language cannot be considered a mere dialect of Herero, as was previously assumed (Estermann 1981). According to this scenario, it is reasonable to assume that the Damara, like the Tjimba, are a cattleless branch of the Himba/Herero who changed their original Herero language after entering into a subordinate, peripatetic-like relationship with the pastoral Nama. Unlike the Damara, the Kuvale share most aspects of their pastoral culture with the Himba and Herero, in spite of their present genetic divergence (Fig. 7).

Recent genome-wide polymorphism data has shown that the Himba, Herero and Damara share a genetic component that is found at lower frequencies in southwestern Bantu populations from the Atlantic coast to the Okavango Delta (Uren et al. 2016; Montinaro et al. 2017). These results, together with our mtDNA and linguistic data, are remarkably consistent with a previously-suggested scenario (Vansina 2004) in which the Bantu pastoralists from Southwest Africa are an offshoot of the Ovambo and/or Nyaneka-Nkhumbi agropastoralists living around the Kunene river basin, who moved into the dry coastal areas of Namibia and Angola. In this framework, it is likely that the different combinations of genetic, linguistic and cultural profiles currently observed in the Himba, Herero, Damara, and Kuvale result from genetic drift, differential admixture and social stratification, instead of reflecting remote geographic origins or assimilation of pre-Bantu components other than “Khoisan” (cf. Vedder and Inskeep, 2003; Möhlig, 2009)

### The peoples of the Kuroca River: Kwisi, Twa and Kwepe

Due to their combination of a peripatetic way of life with a physical appearance that is indistinguishable from their Bantu neighbors, the Kwisi, Twa and Kwepe are frequently seen as the Angolan representatives of a wider group of populations whose origins are often linked to hypothetical pre-Bantu populations different from the Kx’a and Tuu-speaking foragers (Westphal 1963; Cashdan 1986; Barnard 1992; Blench 2006; Güldemann 2008).

While our results show that the Kwisi and the Twa form a relatively homogeneous group that is remarkably different from all other southern African peoples (Figs. 2a and 6a), it is doubtful whether this differentiation could entirely reflect the genetic composition of a pre-Bantu remnant population.

The uniqueness of the two populations can be attributed to their high frequencies of subhaplogroups L0a1b1 (21%), L0a1b2 (11%), L0a2a1b (31%) and L1c1b (22%), which represent approximately 85% on average of their mtDNA composition and are collectively much less frequent in the Himba (18%) and Kuvale (8%) (Fig. 2b; Table S4). Among these four subhaplogroups, L0a1b1 and L0a2a1b are most probably of Bantu origin, since their haplotypes are molecularly close to sequences that are observed in several Bantu-speaking populations from Zambia and Botswana (Fig. 2b; Fig. S5b, d). Haplotypes from subhaplogroups L0a1b2 and L1c1b are confined to the Angolan Namib and have a less clear origin (Fig. S5c, i). While the long internal branches to their closest sequences suggest ancient isolation (Fig. S5c, i), this pattern might also be due to insufficient sampling (Kivisild 2006), or fragmentation of a large ancestral population (Nielsen and Beaumont 2009).

Additional evidence for a link between the Kwisi and Twa and other Bantu peoples of the region is provided by the time depth and genealogical consistency of their clan system (see above), which further suggest that they are likely to be part of the constellation of matriclanic peoples that spread across southwestern Africa (Table S7; Fig. 5b).

In this context, the genetic uniqueness of the Twa/ Kwisi is probably better understood in the frame of a fusion-fission model, where the effects of genetic drift on mtDNA variation are enhanced by the influence of matrilineal kinship on population splitting and ethnogenesis (Neel and Salzano 1967; Fix 1999). Moreover, it is likely that this genetic differentiation was maintained and reinforced by the highly hierarchized social setting of pastoral societies, where impoverished cattleless peoples are marginalized by their dominant neighbors (Vansina 2004).

The relationships between the Twa, Kwisi and Kwepe have also been a matter of contention (Almeida 1965; Estermann 1976; Cashdan 1986). Recently, based on the fact that the Kwepe formerly spoke Kwadi, and on the conclusion that this language could be grouped with Khoe in a single family, Güldemann (2008) suggested that the Kwepe were part of a putative pre-Bantu Khoe-Kwadi migration introducing pastoralism from eastern to southern Africa.

Our results show that the Kwepe have a very homogeneous mtDNA profile (only 5 different haplotypes; Table S4) that bears no resemblance to any other Khoe-Kwadi-speaking population and is largely shared with their neighbors from the Angolan Namib (Fig. S1). While the most common haplotype among the Kwepe is an L3f1b4 lineage (49%) with a likely Himba origin (Fig. 2b; Fig. S1), the other Kwepe haplotypes all belong to subhaplogroups L0a1b1 (27%) or L1c1b (24%) that are more common and diverse in the Twa and Kwisi (Fig. 2b; Fig. S1). These observations suggest that the Kwisi, Twa and Kwepe, who have overlapping residential areas around the Kuroca intermittent river (Fig. 1), were originally the same people, and that the Kwepe mtDNA pool was disproportionally impacted by a single woman, or a kin group, migrating out of the Himba. The genetic similarity of the Kwepe to immediate geographic neighbors displaying Bantu-related mtDNA profiles, rather than to other Khoe-Kwadi-speaking groups, suggests that their former use of Kwadi resulted from language shift after contact with a group of migrants that brought the Kwadi language to the Angolan Namib. So far the only available evidence for a possible genetic contribution of any Khoe-Kwadi migrants to the area is the occurrence in all Namib populations of the lactase persistence −14010^*^C allele (Pinto et al. 2016), which is found with elevated frequencies in several Khoe-Kwadi-speaking peoples of southern Africa (Macholdt et al. 2014). This evidence suggests that there might have been a measurable genetic impact associated with the original Kwadi- speakers that is not captured in the maternal lineages, and might be revealed by Y-chromosome markers and autosomal genome-wide data (currently under analysis).

In any case, the association of the Kwepe with the Kwadi language and a mtDNA profile that is largely derived from the Bantu, combined with the possibility that the Damara represent a branch of the Herero (see above), has important implications for the understanding of the spread of the Khoe-Kwadi family and pastoralism across southern Africa. When linguistically and geographically diverse populations from the region are compared, the most remarkable characteristic of all Khoe-Kwadi speaking peoples is their lack of a common mtDNA genetic heritage (Fig. 6a; Fig. S4). This absence of an mtDNA identity is paralleled by recent data on autosomal DNA variation, showing that many Khoe-Kwadi-speaking groups are genetically closer to populations occupying the same broad geographical area than they are to each other (Pickrell et al. 2012; Uren et al. 2016; Montinaro et al. 2017). Taken together, these patterns suggest that the spread of Khoe-Kwadi and its putative pastoral innovations were part of a complex process that cannot be simply modelled by a wave of advance similar to the spread of agriculture in Europe (Pinhasi et al. 2005), nor by a rapid replacement model, analogous to the Bantu expansions (reviewed in Rocha and Fehn 2016; see also Diamond and Bellwood 2003). It seems more likely that many southern African groups adopted the Khoe-Kwadi language (and occasionally pastoralism) with only a small genetic contribution of incoming Khoe-Kwadi migrants. Our results now indicate that this type of cultural shift not only affected indigenous “Khoisan” foragers, but also impacted Bantu populations from southwestern Africa, leading to the emergence of new ethnic identities that are commonly perceived as enigmatic.

## ACKNOWLEDGEMENTS

The authors thank all sample donors for their participation in this study, the governments of Namibe and Kunene Provinces in Angola for supporting our work, João Guerra, Raimundo Dungulo, and Serafim Nemésio for assistance in the preparation of field work, António Mbeape, José Domingos, and Okongo Toko for assistance with sample collection, Roland Schröder and Enrico Macholdt for assistance in the lab, Shengyu Ni for assistance in the processing of sequencing data, Michael Dannemann, Cesare de Filippo and António M Santos for help in statistical analyses, and Chiara Barbieri for valuable advice. Financial support for this research was provided by FEDER funds through the Operational Programme for Competitiveness Factors— COMPETE, by National Funds through FCT—Foundation for Science and Technology under the PTDC/BIA-EVF/ 2907/2012 and FCOMP-01-0124-FEDER- 028341, and by the Max Planck Society. SO was supported by the FCT grant SFRH/BD/85776/2012, and MG-V by FP7-REGPOT-2011-1-project 286431, UID/BIA/50027/2013 and POCI-01-0145-FEDER-006821. BP acknowledges the LABEX ASLAN (ANR-10-LABX-0081) of Université de Lyon for its financial support within the program “Investissements d’Avenir” (ANR-11-IDEX-0007) of the French government operated by the National Research Agency (ANR). This is scientific paper no. 7 from the Portuguese-Angolan TwinLab established between CIBIO/InBIO and ISCED/Huíla, Lubango. This work is dedicated to the memory of our colleague Samuel Aço.

